# SoyFGB v2.0: a unique access to variations of Chinese Soybean Gene Bank (CNSGB) germplasm

**DOI:** 10.1101/2021.12.28.474253

**Authors:** Tianqing Zheng, Yinghui Li, Yanfei Li, Shengrui Zhang, Chunchao Wang, Fan Zhang, Lina Zhang, Xiangyun Wu, Yu Tian, Shan Jiang, Jianlong Xu, Lijuan Qiu

## Abstract

In Chinese National Soybean GeneBank (CNSGB), we have collected more than 30,000 soybean accessions. However, data sharing for soybean remains an especially ‘sensitive’ question, and how to share the genome variations within rule frame has been bothering the soybean germplasm workers for a long time.

Here we release a big data source named Soybean Functional Genomics & Breeding database (SoyFGB v2.0) (https://sfgb.rmbreeding.cn/), which embed a core collection of 2,214 soybean resequencing genome (2K-SG) from the CNSGB germplasm. This source presents a unique example which may help elucidating the following three major functions for multiple genome data mining with general interests for plant researchers. 1) On-line analysis tools are provided by the ‘Analysis’ module for haplotype mining in high-throughput genotyped germplasms with different methods. 2) Variations for 2K-SG are provided in SoyFGB v2.0 by Browse module which contains two functions of ‘SNP’ and ‘InDel’. Together with the ‘Gene (SNP & InDel)’ function embedded in Search module, the genotypic information of 2K-SG for targeting gene / region is accessible. 3) Scaled phenotype data of 42 traits, including 9 quality and 33 quantitative traits are provided by SoyFGB v2.0. With the scaled-phenotype data search and seed request tools under a control list, the germplasm information could be shared without direct downloading the unpublished phenotypic data or information of sensitive germplasms.

In a word, the mode of data mining and sharing underlies SoyFGB v2.0 may inspire more ideas for works on genome resources of not only soybean but also the other plants.

## Introduction

In an era of genomic information moving forward from theory to application, two key barriers still exist to prevent the widespread sharing of big data: (1) Phenotype is a leading factor for both breeding and genetic analysis. When handling big datasets, deciding how to share germplasms with unpublished phenotypic data, especially quantitative trait data, remains difficult; (2) Genotypic data typically require large storage capacity and relatively infrequent access, whereas germplasm data with phenotypic information require relatively little storage space but frequent access. Thus, how to balance between efficiency and cost is challenging for such databases, especially for researcher-maintained data sources.

Soybean (*Glycine max*) is one of the most important plant sources of protein and oil, and a model crop for legume genome research. Worldwide genebanks such as the Chinese National Soybean GeneBank (CNSGB) and the USDA-ARS Soybean Germplasm Collection contain more than 170,000 soybean accessions, including cultivated soybean (*G. max*) and its progenitor *G. soja*. However, in this era of genomics-based breeding, the sharing of big data, especially the genome sequencing data for multiple soybean accessions remains a bottleneck.

Phytozome (Goodstein et al., 2012) is a popular online resource for plant researchers, and it makes available different versions of reference genomes. Soybase (Grant et al., 2010) provides a unique source of soybean genetic information for multiple soybean genomes based on chip (SoySNP50K) data. In recent years, numerous studies have reported re-sequenced soybean genomes (Li et al., 2014; Liu et al., 2020b; Torkamaneh et al., 2020). The MBKbase (Peng et al., 2020) is also going to release a set of germplasm sequencing data (http://www.mbkbase.org/soybean) based on a recent pan-genome report(Liu et al., 2020a), of which germplasm list for 522 accessions is recently accessible through National Genomic Data Center (Li et al., 2020). LegumeIP, an integrative database for comparative genomics and transcriptomics of model legumes, which has recently been updated to its third version (Dai et al., 2020).

Even so, comparing to other crop species such as rice, access to soybean multiple genome data, especially for haplotype mining with phenotypic data still remains quite limited. Furthermore, soybean is always on the control list for germplasm share. Thus, how to share global soybean collections, especially the core collection with both phenotype and genome data, has become an urgent request by plant researchers.

## Results

In SoyFGB v2.0, we included three major modules, which are Search, Browse, and Analysis. The Search module contains four functions of ‘Germplasm’, ‘Phenotype’, ‘Gene (SNP & InDel)’, and ‘Knowledge’. Users can select favourable germplasms by phenotype scaling in ‘Phenotype’ or by target gene variations embedded in ‘Gene (SNP & InDel)’ module. More information about 2K-SG or soybean was supplied by ‘Germplasm’ and ‘Knowledge’ functions. With the Browse module, the SNP or InDel variations were accessible in a view of genome browser embedded in ‘SNP’ and ‘InDel’ functions, respectively. In the Analysis module, three functions named ‘Hap-GWAS’, ‘Soy_Haplotype’, and ‘Intro_Hap’ were supplied. With these tools, user may carry out a deep mining for haplotypes in genotyped soybean germplasm.

Typical use of SoyFGB v2.0 are demonstrated with followed user cases as example:

### 1) Haplotype mining with embedded/user-owned 2K-SG phenotypic data

In the ‘Soy_Haplotype’ function, user may define a target region with gene name, physical range or a set of SNPs, to mine the possible haplotype variations from the 2K-SG. With the embedded or user input phenotypic data, the mean values of target traits for different haplotypes are available. The possible donor lists are provided with a straight-forward statistical analysis based on ANOVA protected t-test as reference for users.

The candidate genes for isoflavone content in soybean were identified by a joint work of bulk segregant analysis (BSA) with a natural population and weighted gene co-expression network analysis (WGCNA) using the transcriptome of different seed development stages. SoyFGB v2.0 provided haplotype analysis function for the candidate genes. Firstly, locus number of one candidate gene ID and the phenotypic data of isoflavone content of 2K-SG from user were submitted to the ‘Soy_Haplotype’ function embedded in Analysis module of SoyFGB v2.0. Then, all the haplotypes of this gene were presented. Subsequently, With the aid of straight-forward statistical analysis between different haplotypes, germplasms harbouring different haplotypes were found to be significantly distinct from each other in isoflavone content. This implies the possible contribution of the candidate gene in regulating isoflavone content of soybean grain. Finally, the haplotype variations including and the germplasm list for the candidate gene were also downloaded for further lab works (Figure 1).

**Figure 1.**
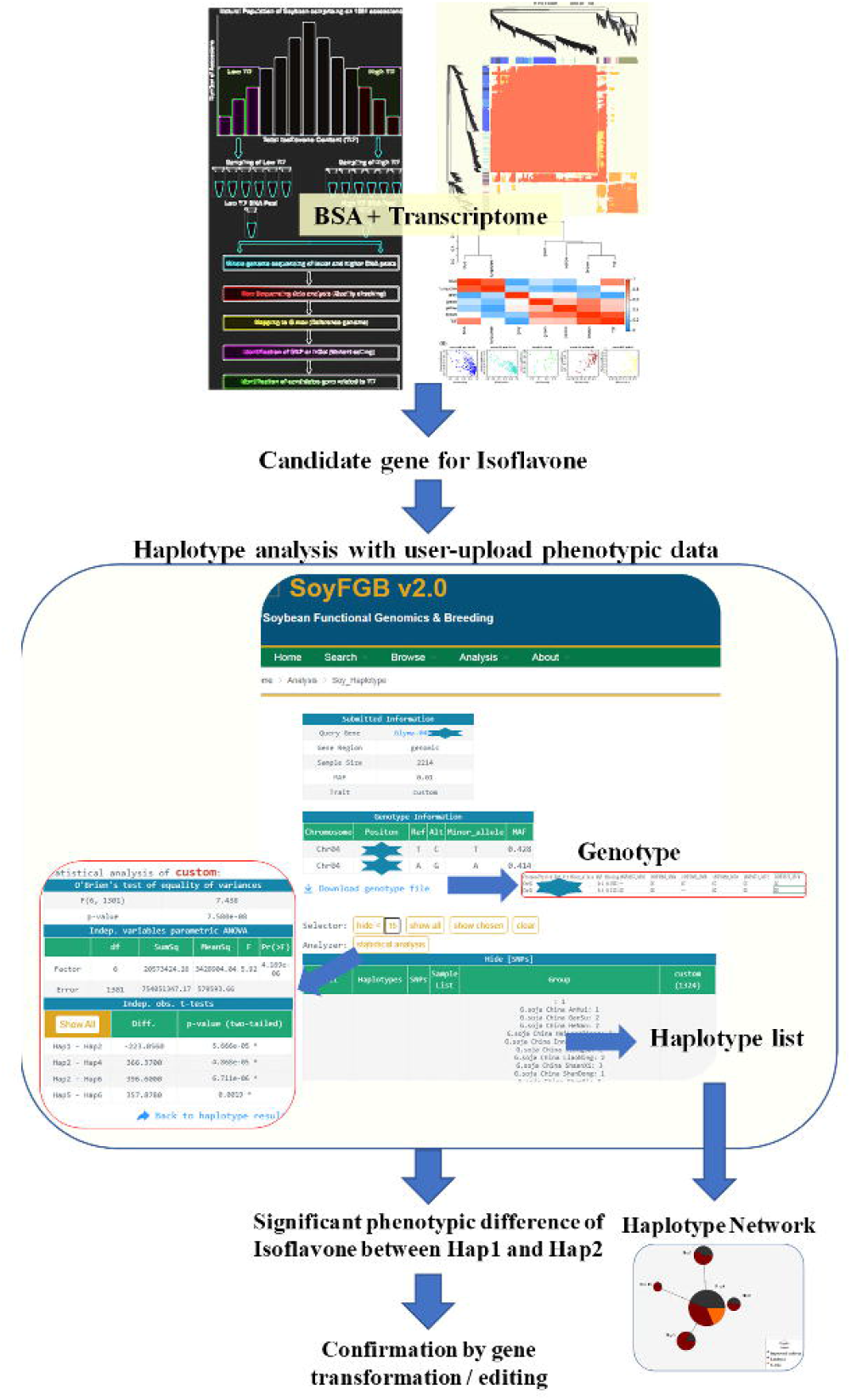
User case of ‘Soy_Haplotype’ function embedded Analysis module of SoyFGB v2.0.

### 2) Finding candidate gene / region with Hap-GWAS function

As shown in Figure 2, an enhanced correlation between the phenotypes and haplotypes would be mined with Hap-GWAS function, which adopted the methodology raised recently (Zhang et al., 2021). In order to save the possible waiting time for this analysis, an email-remind system was adopted. Once the analysis results were ready, a remind-email with a direct access link to the output would be sent to the mailbox defined by user. Together with the instant screening with the previous ‘Soy_Haplotype’ function, correlations between the phenotype and haplotypes may be fully mined at different depths.

**Figure 2.**
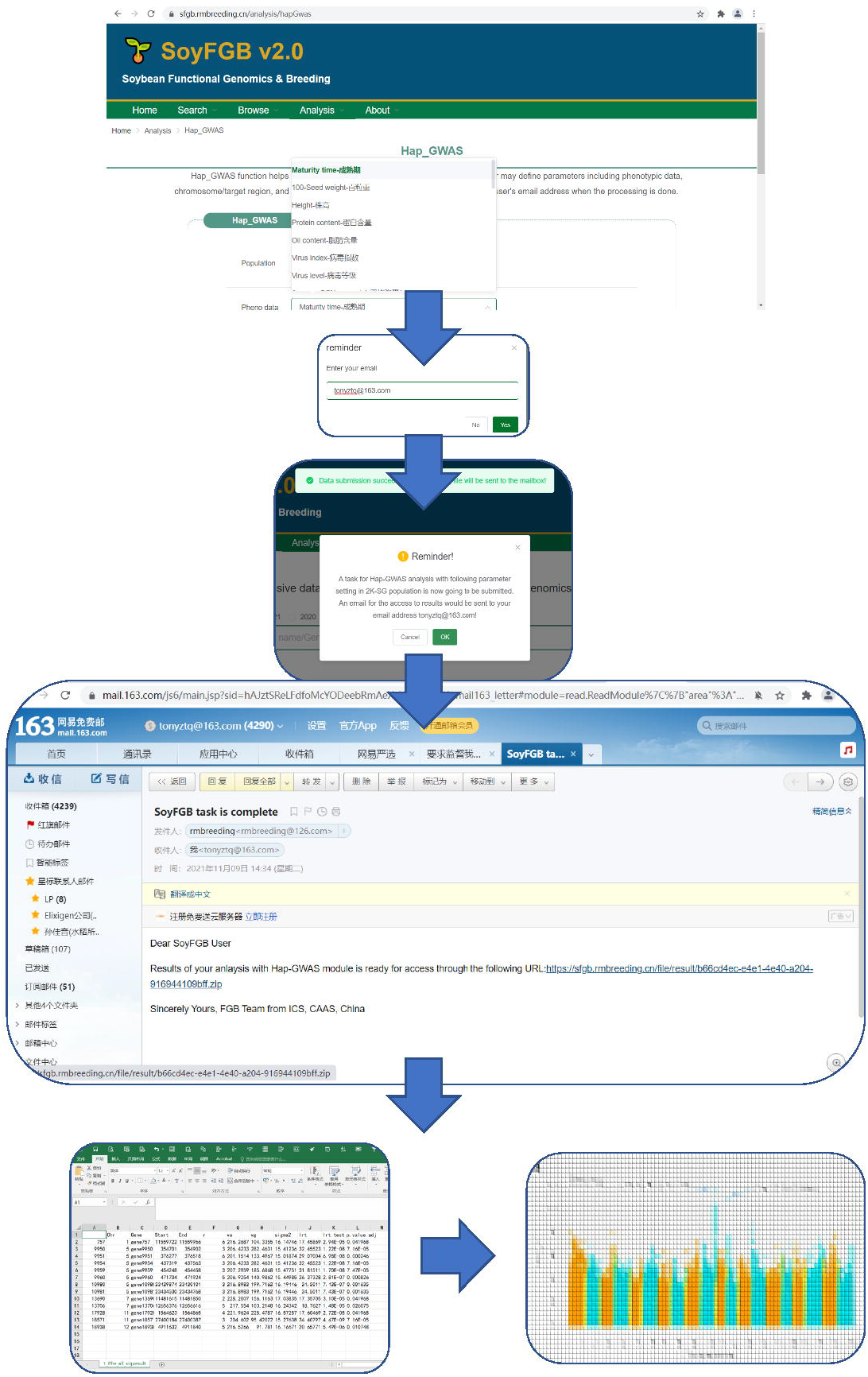
Find target gene / region with ‘Hap-GWAS’ function embedded in Analysis module of SoyFGB v2.0.

### 3) Intro_Hap function to mine haplotypes with SNP chip data

Considering more and more data sets accumulated by relatively lower density genotyping methods, e.g. SNP chip, an analysis tool for haplotype in target region using this type of data were also provided with ‘Intro_Hap’ function for both populations with/without known parents. Since the request for this module is recently raised by users during out indoor testing period, it is still looking forward to more users’ responses within a 6-month open period since this release of SoyFGB v2.0.

### 4) Exploring germplasms based on scaled-phenotype or accession information

A typical pre-breeding / forward genetics scheme starts with phenotyping. In SoyFGB v2.0, a set of scaled data covering 42 traits, including 9 quality and 33 quantitative traits (maturity time, 100-seed weight, height, protein content, oil content, virus index, virus level, average SCN amount 1, SCN infection level 1, average SCN amount 2, SCN infection level 2, average SCN amount 3, SCN infection level 3, average SCN amount 4, SCN infection level 4, average SCN amount 5, SCN infection level 5, cystine acid content, methionine acid content, stearic acid content, palmitic acid content, oleic acid content, linoleic acid content, linolenic acid content, salt tolerance during germination, salt tolerance of seedlings, sprouting drought tolerance, drought tolerance of adult plants, cold tolerance during germination, cold tolerance of seedlings, resistance to soybean rust, type of reaction to soybean rust, soybean rust infection level) were embedded in Search based on ‘Phenotype’ function. The user can screen 2K-SG germplasm with scaled phenotypic data. A three-step route can be followed for exploring elite donors for a breeding scheme: (a) target trait scaling, demonstrated herein by screening the top 30% protein content as an example, which includes 13 samples; (b) from these samples, favourable early mature (top 50%) samples were further screened, and favourable samples may be added to create a list of candidate germplasms (three samples); (c) the user can then export a list of candidate donors for different breeding schemes based on the two grouping levels. An easy way to ‘Seed Request’ function is available through just one click on a key named ‘Request Germplasm’.

### 5) Searching for variation within candidate genes for favourable germplasm

A route for shortlisting candidate genes using SoyFGB v2.0 is shown in Figure 3, and involves the following: (a) get a target region by mapping methods such as GWAS or sorting accessions with favourable target traits, (b) using ‘SNP’ or ‘InDel’ functions in Browse module, variations within target gene / region can be explored, (c) inputting target region or gene locus ID into the ‘Gene(SNP/InDel)’ function of Search module, (d) with the genotype information (SNP or InDel) downloaded, users can carry out further work with primer design and wet-lab confirm.

**Figure 3.**
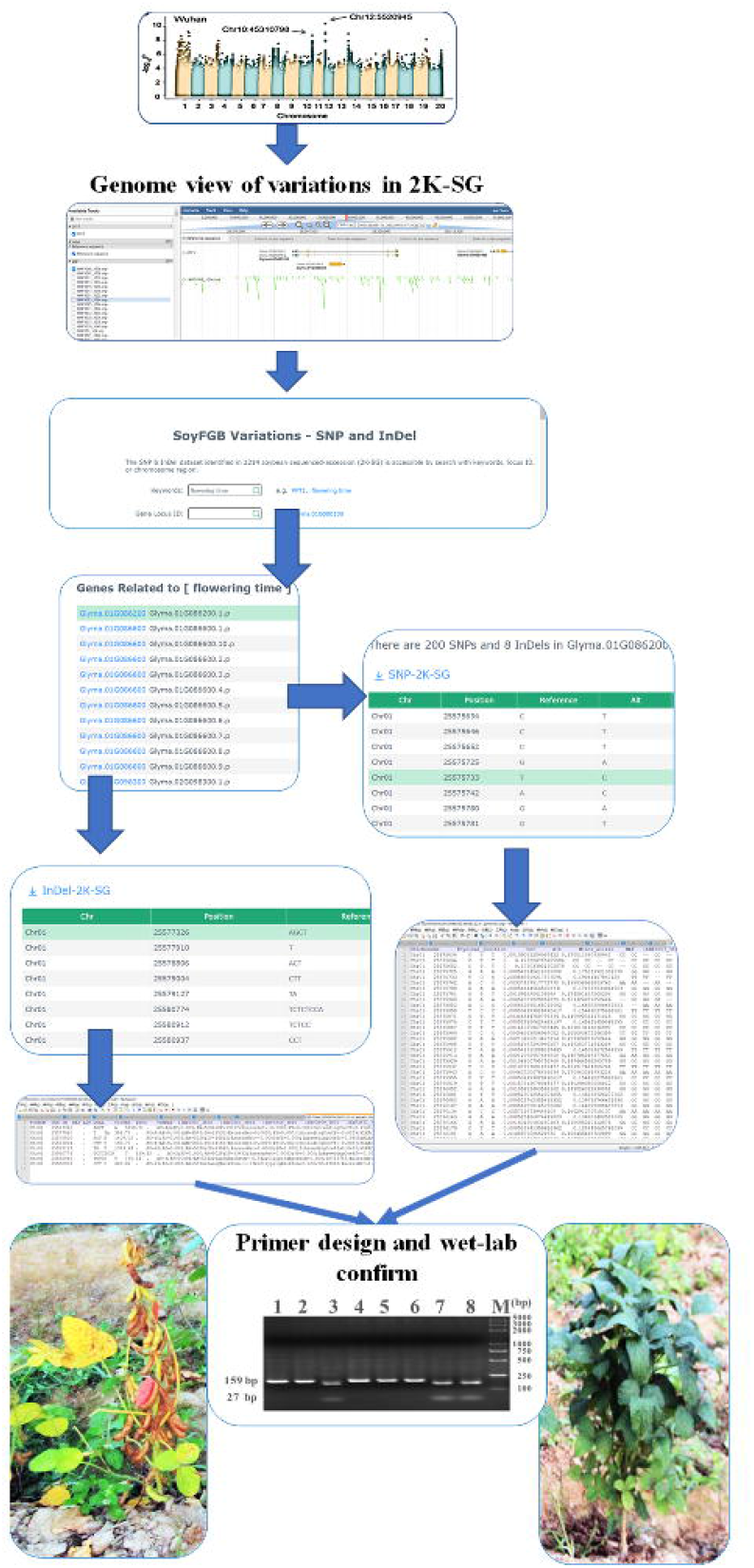
User case of variation mining for flowering time with Search module of SoyFGB v2.0.

## Discussion

SoyFGB v2.0 is accessible through the following URL: https://sfgb.rmbreeding.cn/. It is designed to be adaptive and responsive to the overwhelming quantity of genomic data and phenotypic data resulting from functional genomic breeding materials, including 2K-SG data. The FGB data sharing mode in SoyFGB v2.0 has characteristics as followings:

(i) Instead of providing a direct download link to raw phenotypic data, a scaled-based phenotypic data-led germplasm sharing mode is employed by SoyFGB v2.0. On the other hand, the correlations between phenotypic and genotypic data, such as GWAS results are commonly provided by search function of other websites (Li et al., 2020; Zhao et al., 2021). SoyFGB v2.0 make an attempt to the online analysis with ‘Hap-GWAS’, ‘Soy_Haplotype’, and ‘Intro_Hap’. Mining elite donors with favourable haplotype are of high value for breeders. This has provided a model which is more conducive to data contributors sharing their own unpublished data with public users.

(ii) To keep up with the development of multiple-client ends, a development framework different from previous FGB website (Wang et al., 2020) has been employed in SoyFGB v2.0. The front end was developed by the Vue-Element-Axios tool, and the back end of was developed by using the java-based Spring Boot tool. The website is driven by Nginx. The RESTful API facilitates easy data access through different client platforms. Additionally, in order to balance efficiency and cost, a distributed database structure was designed for SoyFGB v2.0 (Figure 4). Phenotypic and genotypic data are stored in servers with different capacities. Phenotypic data are managed by an instant-response server with a relatively small storage capacity using the MySQL database. Genotypic data are stored in a server with a slower response but a larger storage capacity. All these attempts are going to meet the possible developing trends including decentralization and multiple accessing ways to biological data in near future.

**Figure 4.**
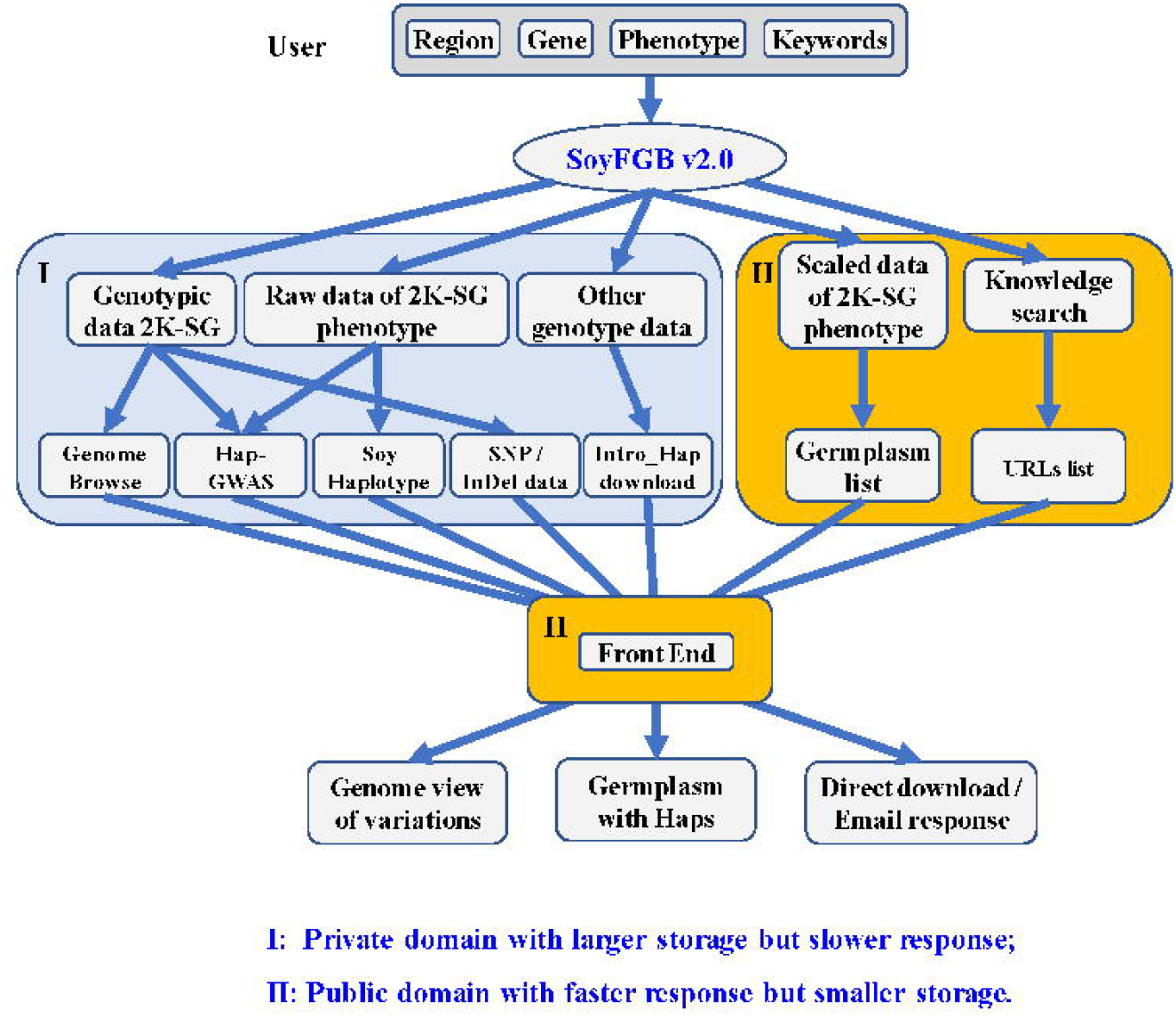
Database structure of SoyFGB v2.0.

(iii) Searching plant omics databases for functional information has grown in popularity (Gui et al., 2020). Accordingly, in SoyFGB v2.0 the Search function is important for helping users to search for useful information inside and/or outside SoyFGB. With all three major modules embedded, users can access 2K-SG data in an effective and efficient manner.

In summary, SoyFGB v2.0 provides a unique example of platform for sharing big datasets (both phenotypic, genotypic, and mining data) from of multiple soybean sequenced accessions. With the development of SoyFGB, the FGB has now evolved into a phenotype-led re-sequencing data sharing mode. This may inspire new ideas for mining complex traits in soybean and other plants. With more and more 2K-SG data published by users who query and use this resources, SoyFGB v2.0 will continue to acquire more and more data, especially phenotypic data, that will be available for users. In future updates, we will focus on data integrating with more dimensions (for both genotypic and phenotypic data) to collate variations in 2K-SG germplasm data from different groups in China and across the world. With progress in genotypic data analysis and phenotypic data collection from re-sequenced genomes, more open links between these datasets is hopefully going to be constructed in progressing versions of SoyFGB. Further development of the FGB and other databases will facilitate a unique access to soybean and other plant collections in genebanks.

## Materials and Methods

As shown in Figure 5, SoyFGB v2.0 includes 2,214 accessions (2K-SG) from four major soybean production and distribution areas (Asia, America, Europe and Africa) based on the core collection strategy of the CNSGB. The 2K-SG dataset comprises four major classes of soybean species; cultivated species (1,993 *G. max*), annual wild species (218 *G. soja*), perennial wild species (2 *G. tomentella*), and others (1 *G. tabacine*). Among them, *G. tomentella* and *G. tabacine* are the only two perennial wild species found in China. Of the 218 *G. soja* accessions, 99.5% were collected from native sources (East Asia). This includes China (179), Korea (10), Japan (19) and Russia (9), providing broad species diversity. Among the 1,993 *G. max* accessions, more than half (56.7%) are landraces primarily collected from China and applied to core collections to represent the diversity of the 23,587 cultivated soybean accessions preserved in the CNSGB. The 862 improved cultivars were collected from 17 countries, especially major soybean producing countries including the USA and China.

**Figure 5.**
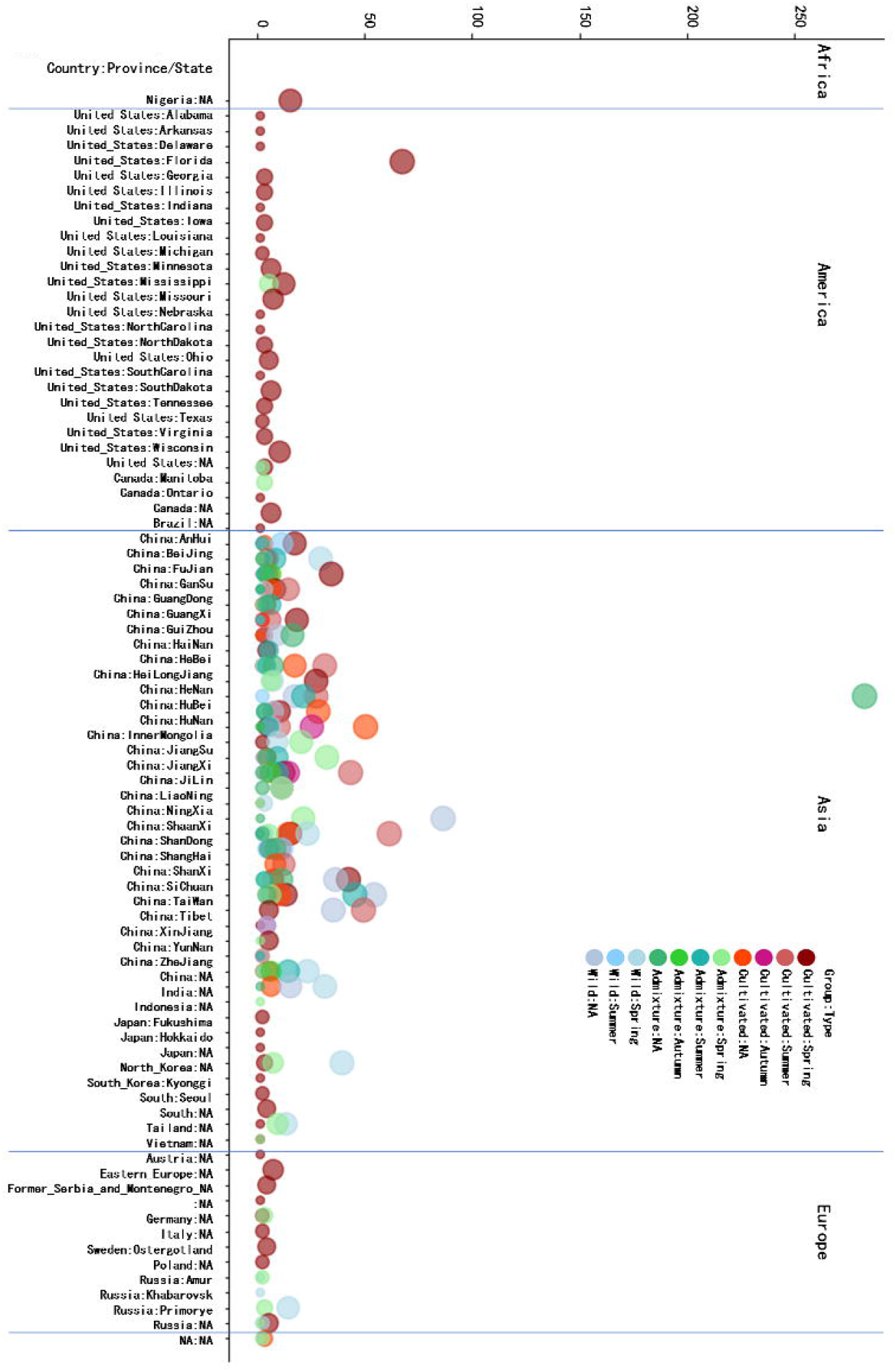
The 2,214 sequenced soybean genome (2K-SG) germplasm embedded in SoyFGB v2.0.

Whole-genome resequencing was carried out according to a standard procedure. Specifically, a genomic library was constructed using a TruseqNano DNA HT sample preparation Kit (Illumina, San Diego, California, USA), purified using an AMPure XP bead system (Beckman Coulter, Brea, California, USA), and the size distribution was analysed by an Agilent 2100 Bioanalyzer (Agilent Technologies, Santa Clara, California, USA) and quantified using real-time PCR. An Illumina Hiseq X platform was then employed to generate ∼10.58 Tb of raw sequences with a read length of 150 bp. Removal of low-quality paired reads resulted in 16.41 Tb of high-quality paired-end reads, including 96.05% with phred quality ≥Q20 and 90.98% with phred quality ≥Q30. The obtained fastq files were submitted to a pipeline composed by BWA (v. 0.7.17-r1188), SAMtools (v.1.39), Sambamba (v.0.6.8), picard (v.2.18.15, http://broadinstitute.github.io/picard), and GATK (version v4.1.2.0) and screened against Williams 82 assembly V2.0 (http://www.phytozome.net/soybean). This yielded 65,374,688 single-nucleotide polymorphisms (SNPs; 60,153,828 bi-allelic) and 10,952,749 InDels (8,349,613 small insertions and deletions <15 bp and <50% missing). Using a standard SNP screening procedure, 8,785,134 highly-credible biallelic SNPs were obtained.

Based on this set of SNPs, two different levels of grouping were carried out and presented in SoyFGB v2.0. Level one (Group 1), the SNP-only level, includes 1507 cultivated, 313 wild, and 394 admixture accessions. Level two (Group 2), based on the output of a two-step grouping, includes a CGCC-based subgrouping and an SNP-based subgrouping within each group. In Group 2, the cultivar group was divided into five subgroups; the Chinese Southern Region (C_SR), the Chinese central region surrounding the middle area downstream of the Yellow River valley (C_CR), the Chinese northern region plus Japan, the Korean peninsula and the Russian far east region (C_NR), America (C_Am), and admixture (C_AD) subgroups. The wild group was divided into four subgroups; the Chinese southern region (W_SR), the Chinese central region surrounding the middle area downstream of the Yellow River valley (W_CR), the Chinese northern region plus Japan, the Korean peninsula and the Russian far east region (W_NR), and admixture (W_AD) subgroups.

It’s known that, InDel and SNP tend to be gathering throughout the genome (Montgomery et al., 2013). Thus, in present release of haplotype analysis, SNP is still the main points, the InDel presented by “-” was also taken into consideration during the analysis. Additionally, heterozygotes were regarded as one type. In the future versions, analysis with models considering more variations including InDel and more reference genomes would be taken into accounts. In ‘Soy_Haplotype’ function, a straight-forward statistical analysis based on ANOVA protected t-test is provided. In the ‘Hap_GWAS’ function, a linear model for GWAS (Zhang et al., 2021) was adopted.

Phenotypic data were obtained from CNSGB accumulated data. Quantitative data were then transformed into scaled form.

## Author Contributions

QLJ, YHL and TQZ conceived the study and drafted the manuscript. YFL, SRZ, YHL, YT, SJ and TQZ contributed to data sharing. CCW and FZ contributed to raw code. LNZ and XYW contributed to database maintenance. JLX contributed to writing and revision.

## Acknowledgements

This work was partially supported by the National Key Research and Development Plan (2016YFD0100201) from MOST, the National Natural Science Foundation of China (31871715), the Central Public-interest Scientific Institution Basal Research Fund (Y2020PT24) and the Agricultural Science and Technology Innovation Program (CAASTIPS, Y2020YJ09) from the Chinese Academy of Agricultural Sciences, the Phenomics project from the Institute of Crop Sciences (ICS2020YJ07BX), and the ‘Green Super Rice’ project from the Bill & Melinda Gates Foundation (OPP1130530).

